# Two loci single particle trajectories analysis: constructing a first passage time statistics of local chromatin exploration

**DOI:** 10.1101/135012

**Authors:** O. Shukron, MH. Hauer, D. Holcman

## Abstract

Stochastic single particle trajectories are used to explore the local chromatin organization. We present here a statistical analysis of the first contact time distributions between two tagged loci recorded experimentally. First, we extract the association and dissociation times from data for various genomic distances between loci and we show that the looping time occurs in confined nanometer regions. Second, we characterize the looping time distribution for two loci in the presence of multiple DNA damages. Finally, we construct a polymer model that accounts for the local chromatin organization before and after a double-stranded DNA break (DSB) to estimate the level of chromatin decompaction. This novel passage time statistics method allows extracting transient dynamic at scales from one to few hundreds of nanometers, predicts the local changes in the number of binding molecules following DSB and can be used to better characterize the local dynamic of the chromatin.

## 1 Author summary

Extracting local properties of the chromatin from temporal tracking of a tagged genomic section remains a challenging task. In this work, we analyze empirical trajectories of two tagged loci, simultaneously tracked over time and positioned at various genomic distance apart. The correlated motion of two loci provides complementary information compared to single ones, contained in the distribution of their recurrence encounter times. We show here how to construct such distribution from empirical data. In particular, We find that this distribution is affected after the induction of damages by Zeocin drugs (causing double strand DNA breaks) and report the changes in the local chromatin environment exploration. Finally, we examine the local chromatin remodeling following a double strand breaks, by using simulation of a randomly cross-linked polymer model. We show that only a small percentage of cross-linker molecules are removed following damages. The analysis presented in this work could be extended to multiple tagged loci.

## 2 Introduction

Analysis of single particle trajectories (SPTs) of a tagged single locus revealed that chromatin dynamics is mostly driven by stochastic forces [1, 2]. The statistic of a locus motion has been characterized as sub-diffusive [3, 4, 6, 7, 8] and confined into nano-domains. The confinement is probably due to an ensemble of local tethering forces generated either at the nuclear periphery [9], or internally [10], where binding molecules such as CTCF or cohesin play a key role [11, 12]. Chromatin dynamics is involved in short-range loop formation in the sub-Mbp scale, and contributes to processes such as gene regulation. However, the analysis of the chromatin dynamics in the sub-Mbp scale is insufficient to describe processes involving long-range chromatin looping (above Mps scale), such as in homologous dsDNA repair. When two neighboring loci located on the same chromosome arm are tracked simultaneously over time, their correlated position can be used to explore the local chromatin organization in the range of tens to hundreds of nanometers (of the order of the genomic distance between the loci).

Statistical parameters characterizing short-range chromatin motion have been studied in stochastic polymer models, starting with the Rouse polymer [13], copolymers [14], the beta polymer [15], and polymer models with additional diffusing or fixed binding molecules [16, 17, 18, 19]. The extracted statistical parameters are the diffusion coefficient, local tethering forces, the radius of gyration, radius of confinement [1, 2], and the distribution of anomalous exponents of tagged loci along the chromatin, which characterizes the deviation from pure diffusion [19, 4].

Here we analyze the transient statistics of two loci SPTs and use it to explore the local chromatin reorganization following DSB and its confining geometry. Thus, we further contribute to the global chromatin reorganization explored in [1]. We adopt here the formalism of Brownian polymer dynamics, as we have already shown [10] that the auto-correlation function of a single locus decays exponentially, but not as power laws, as would be predicted by the fractional Brownian motion description [5]. Specifically, we explore the chromatin state from the transient statistics of recurrent visits of two tagged loci. This approach is new and is not contained in other work involving two spots trajectories, which use equilibrium thermodynamic models for steady-state encounter frequency [20] or specific chromatin arrangement [21]. We study the distributions of 1) the first encounter time (FET) and 2) the first dissociation time (FDT) of two tagged loci. The FET is defined as the first arrival time of one locus to the neighborhood of the second, while the FDT is the first time the two loci are separated by a given distance. The statistics of FDT and FET is not contained in moments associated with each locus separately, but revealed by their correlated motion.

This article is organized as follow: in the first part, we introduce and estimate the FET and FDT distribution from SPTs of two loci (data from [22]). In the second part, we analyze empirical data of loci motion before and after the induction of DNA damages by Zeocin (data from [1]). The local effects of DSBs on the loci motion was not the goal in [1], but multiple DSBs and single stand breaks (caused by Zeocin), together with a strong DNA damage checkpoint response can trigger global chromatin changes. We shall study here the consequences of multiple tether losses on the chromatin not just around the break site, on the local loci motion. In the third part, we use a randomly cross-linked (RCL) polymer model [17, 19] to simulate the trajectories of two loci following a DSB on the DNA strand between them and evaluate the number of binding molecules required to restrict their motion. We thus use the RCL polymer to explore the chromatin reorganization on the scale of a single DSB. In the last section, we estimate the number of binding molecules required to obtain SPTs with the same statistics as the measured ones. We conclude that the statistics of two correlated loci provide complementary information about local chromatin organization, not contained in the statistics of individual non-correlated loci. The present method is general and can be applied to any SPT of any number of loci. It can further be used to reveal characteristic lengths, local chromatin dynamics, remodeling following DSB and estimate the changes in the number of molecular interactions.

## 3 Results

### 3.1 First passage time analysis

The construction of the present statistical method is based on the first passage time for two loci entering and exiting a small ball of radius *∊* (that can vary continuously). We will thus estimate the FET and FDT (introduced above). The statistics of these times contain information about the local chromatin organization at a scale of one to few hundreds of nanometers, because the fluctuations in loci distance depend not only on their stochastic dynamics but also on the restricted geometry. We now briefly recall the published data we will used to construct the analysis. In the data of [22], two fluorescently tagged loci are tracked over a course of 60-120s. We only used recording for which the time interval did not exceed 1s. The experiment is repeated for seven DNA strains of genomic length between the tagged loci between 25-100 kbp. We also use the dataset reported in [1], which tracks two tagged loci located on yeast chromosome III, at a genomic distance of 50 kbp, at time intervals of 300ms for a total of 60s. The trajectories of two loci are tracked after the induction of DSB breaks uniformly over the genome by Zeocin 500 *μg/ml*.

We first analyze trajectories of two tagged loci of [22], when they are separated by various genomic distances: Δ = 25.3; 42.3; 51.3; 71, and 100.8 *kbp*. The distance *d*(*t*) = *dist*(*X*(*t*), *Y* (*t*)) between the two trajectories *X*(*t*) and *Y* (*t*) fluctuates in time, thus we estimate the distribution of the FET _*τE*_ and the FDT _*τD*_ (Fig. 1A). The FET is the first time the distance between the two loci becomes less than ∊, when the initial distance is larger. The FDT is defined as the first time that the distance between the two loci reaches ∊ when they are initially inside a ball of radius ∊. The FET (FDT) are collected between successive dissociation (association) events, after which we reset the time to *t* = 0: by definition:

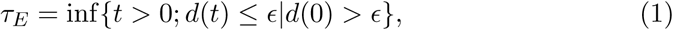

and

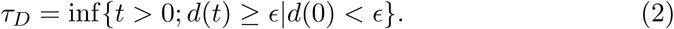

**Figure 1:**
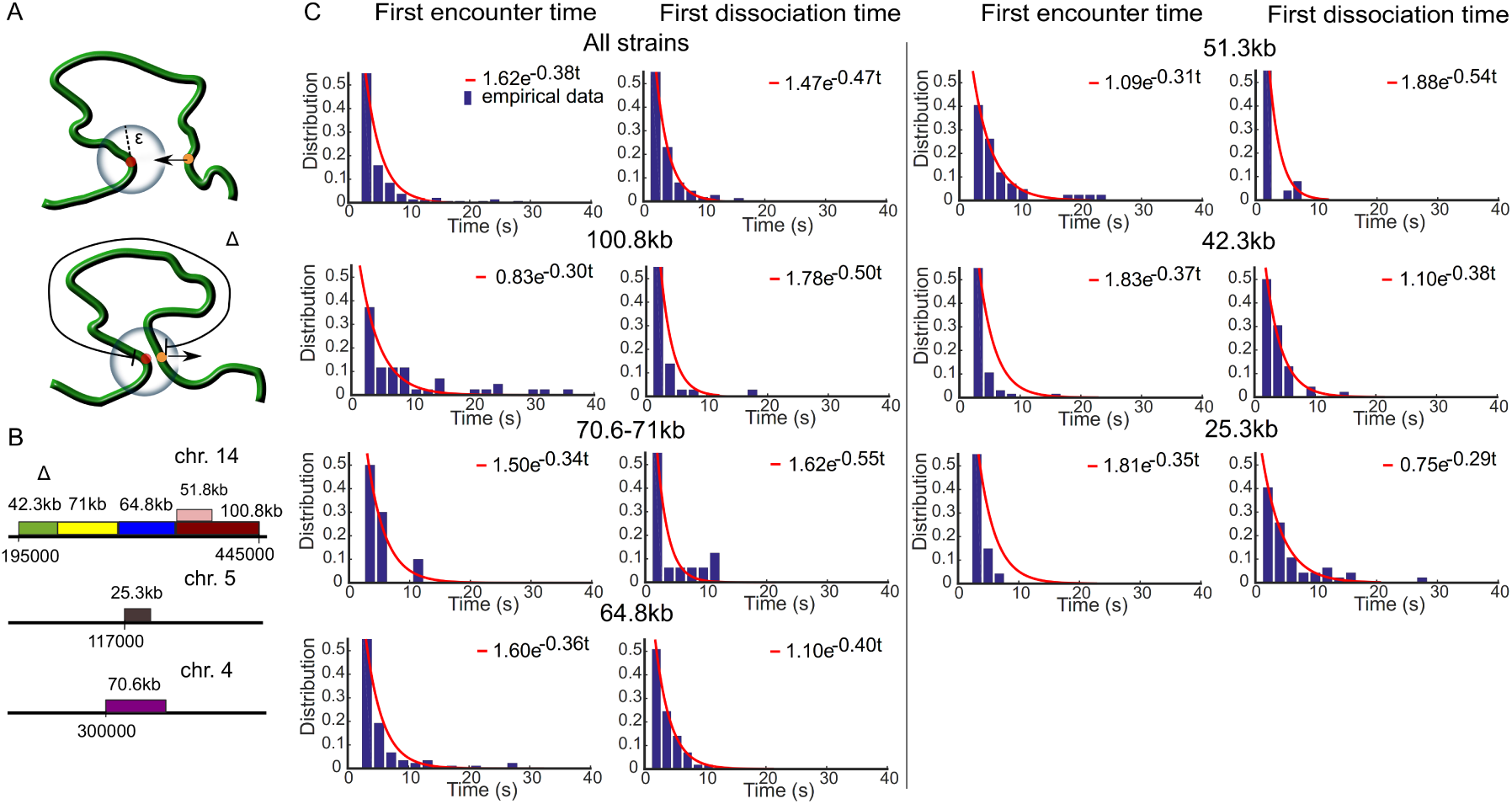
Statistics of two loci trajectories. **A.** schematic representation of the first encounter time (FET) *τ*_*E*_ (upper) and the first dissociation time (FDT) *τ*_*D*_ (lower). The FET is computed when the two loci are within an encounter distance ∊when they are initially apart. The FDT is computed when the distance between two loci is larger than *∊* when they initially encountered. The genomic distances are Δ ∈ [25.3, 100.8] kbp between tagged loci **B.** Experimental setting for tagging seven chromatin strains by inserting lac and tet flanking operators at their ends on chromosome 4,5 and 14 [22] **C.** distribution of the FET (left column) and the FDT (right column) with respect to Δ, fitted with *a* exp(−*λt*), with *a* a constant.

In practice, we constructed the distributions of *τ*_*E*_, *τ*_*D*_ for a continuum of encounter distances *∊* that varies in the range 150-500 nm.

We gathered the distributions of the FET and FDT for seven various DNA strains of genomic length Δ = 25−108 kbp [22] (Fig. 1B). As predicted by the polymer looping theory in confined domains [23, 15] (formula 5 of the Method), the distribution (Fig. 1C red curves) follows a single exponential decay, with rate *λ*, which is the reciprocal of the mean FET (MFET) between the two loci. Using an exponential fit to the data for all strains of length Δ, we find that the MFET slightly decreases from 3.2 s for Δ = 25 kbp to 2 s for Δ = 108 kbp (Fig. 2A blue circles).

**Figure 2:**
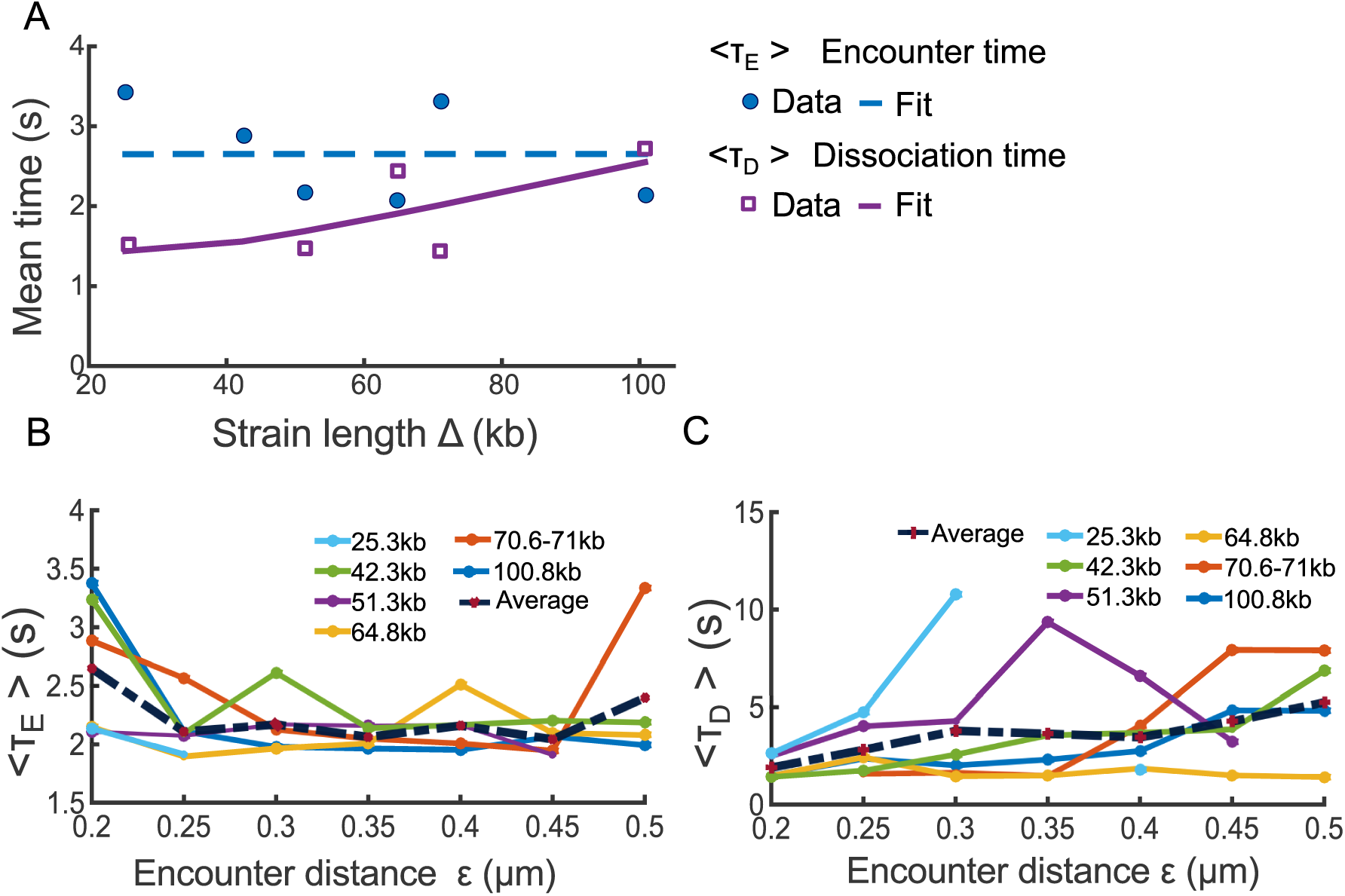
Effect of the genomic separation distance Δ and the encounter distance ∊. **A.** The mean first encounter time (MFET) data (blue circles) are fitted using eq. 6 (blue dashed). The mean first dissociation time (MFDT) data (purple squares) is fitted using eq. 9 *a*_2_Δ exp(*b*_2_*/*Δ) (purple curve), where *a*_2_ = 0.01, *b*_2_ = 40.36 **B.** The MFET 〈*τ*_*E*_〉 for 6 strain lengths Δ kbp, shown in Fig. 1B, where the encounter distance *∊* varies in 0.2 and *μ*m. **C.** MFDT 〈*τ*_*D*_〉 extracted from Fig. 1B and plotted for all Δ with respect to the encounter distance ∊.

To estimate the effect of chromatin confinement on transient properties, we used the formula 6, derived for confined polymer, to fit the MFET of the two loci for all Δ. Because the two loci are located along the same chromosome arm [22], we model them as the two end monomers of a polymer chains with *N* monomers. To fit the the MFET data using 6, we use Δ ∈ [25, 108] kbp, *b* = 0.2 *μ*m, the length of 1 bp to be 3 x 10^−4^ *μ*m, the number of monomers *N* = (3 × 10^−4^ Δ/*b* and the parameters *D* = 8 x 10^−3^ *μm*^2^*/s*, *κ* = 1.75 x 10^−2^ N/m and ∊ = 0.2 *μ*m (see Table 1). We find the value for the confined parameters *β* = 2.4 *μm*^−2^, and substituting in 7, we finally obtain the radius of confinement of *A* = 0.5 *μ*m in agreement with data presented in [22]. Furthermore, the MFET in a confined environment does not exceed a limit of 2.65 s for all genomic distances between tagged loci (Fig. 2A dashed blue), suggesting that the dynamics has already reached the asymptotic limit and thus the loci are confined at all scales. We conclude that the motion of two loci located in the range 25-108 kbp is largely influenced by the local chromatin confinement.

**Table 1:** values of simulation parameters

| Parameter | Value | Description |
| --- | --- | --- |
| N | 100 | number of monomers |
| b | 200 nm | STD of adjacent monomers distance |
| D | $8 \times 10^{-3} \mu m^2/s$ | Diffusion coefficient [15] |
| $\epsilon$ | 0.2 $\mu m$ | Encounter distance |
| $\kappa$ | $1.75 \times 10^{-2}$ N/m | Spring constant[34] |
| $\gamma$ | $3.1 \times 10^{-4}$ Ns/m | friction coefficient [34] |

The stochastic model for the FDT is the escape problem from a parabolic potential well [31]. The mean escape time is given by formula 9 (Methods), which shows that the dissociation time increases with the genomic length Δ. We thus fitted the mean FDT (MFET) data points using formula 9 (Fig. 2A, purple squares), confirming that the MFDT increases from 1.5 s to 2.5 s when Δ increases from 25 to 108 kbps (Fig. 2A) (Using a Matlab fit, we estimated the parameters of relation 9 to be *a*_2_ = 0.01, *b*_2_ = 40.36).

To evaluate the sensitivity of our approach to the choice of encounter distance ∊, we estimated the FET and FDT when ∊ varied in the range 0.2 − 0.5 *μ*m. For ∊ ∈ 0.2 − 0.25 *μ*m, the MFET decreased from 2.7 s for ∊ = 0.2 *μ*m to a 2.1 s when ∊ = 0.25 *μ*m (Fig. 2B dashed). In the range ∊ ∈ [0.25, 0.5] *μ*m, we find that the MFET is almost constant, independent of ∊ with an average of 2.1 s (Fig. 2B left). This result shows that it takes around 2 seconds for the two loci to meet and thus to explore a ball of radius of 0.25 *μ*m. Indeed, the MFET is the time to meet when almost all points of the domain have been visited [24].

We conclude that any loci explore constantly and recurrently the neighborhood of the chromatin with a time constant of 2 seconds in a tubular neighborhood of 0.25 *μ*m and this spatial constraint does not depend on the position of the locus. We note that this result about the exploration and the recurrence exploration is not contained in the statistics SPT of a single loci, because a reference point is needed. Finally, we find that the MFDT increases with ∊ for all DNA strain of length Δ (Fig. 2C), from an average of 2 s for ∊ = 0.2 *μm* to 5 s when ∊ = 0.5 *μ*m.

### 3.2 Loci dynamics in the presence of double-strand DNA break

To continue exploring how two loci trajectories provide information about local chromatin organization, we focus now on the consequences of double-strand DNA breaks (DSBs) on chromatin dynamic. For that purpose, we analyzed the transient statistics of two loci before and after treatment with the radiomimetic drug Zeocin (data presented in [1]). The Zeocin drug induces uniformly distributed DSBs on the chromatin, leading to chromatin expansion and enhanced chromatin flexibility [1]. Thus, we repeated the FET and FDT statistical analysis (Fig. 3A) as described in the previous subsection. As predicted by the polymer model theory, the FET and FDT follow a Poisson distribution and we fitted a single exponential (formula 5) to the empirical distributions (Fig. 3B). We then computed the MFET and MFDT for encounter distances *∊* in the range 0.1 − 0.5 *μ*m.

**Figure 3:**
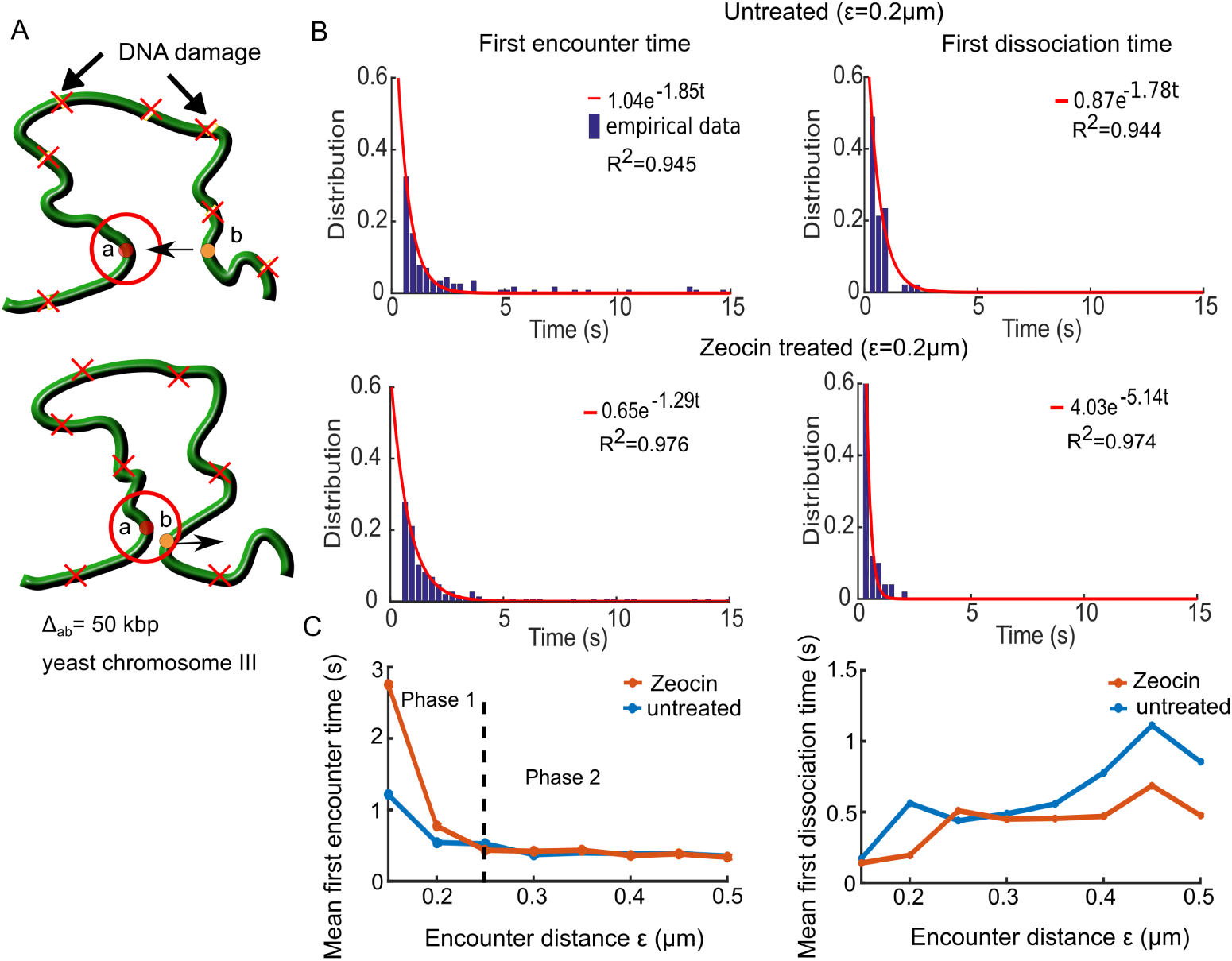
Two loci dynamics before and after Zeocin treatment. **A.** Two tagged loci (a and b, circles) separated by a genomic distance Δ_*ab*_ = 50 kbp. When the loci are within a distance ∊ *μ*m (red circle), they are considered to encounter for computing the FET, and above ∊ (lower) for the FDT. We analyzed the untreated and Zeocin treated cases, where Zeocin induces DNA damages (red X) at random positions along the DNA. **B** the FET (left column) and FDT (right) empirical distributions in the untreated (upper) and Zeocin treated (lower) cases, fitted by *a* exp(−*λt*) (red curves), where the reciprocal of *λ* is the MFET and MFDT in their respective cases. *R*^2^ values from the fit are reported in each box **C** The MFET (left) is plotted with respect to the encounter distance *E* for the untreated (blue) and Zeocin treated (orange) cases. For the MFET (left), both curves are at a plateau of at 0.5 s (phase 2) above ∊ *>* 0.25 *μ*m. The MFDT (right) increase with ∊ from 0.2 s at *∊* = 0.15 *μ*m to 0.5 s at ∊ = 0.5 *μ*m for the untreated (blue) and Zeocin treated (orange) case.

For both the untreated and Zeocin treated data, the MFET graphs Fig. 3C show two phases: in the first, when ∊ ∈ [0, 0.2] *μ*m, the MFET decays with the radius ∊, while in the second phase (∊ ∈ [0.2, 0.5] *μ*m), it is independent (Fig. 3C). The boundary between the two phases at ∊ = 0.25 *μ*m indicates that this length is a characteristic of local chromatin folding and crowding. Interestingly, following Zeocin treatment, the MFET increases at a scale lower than 0.25 *µ*m, compared to the untreated case, probably due to the local chromatin expansion around DSBs. Furthermore the increase of the confinement length *L*_*c*_, when the chromatin is decompacted [1] and the restriction of the loci dynamics can be due to repair molecules.

To further investigate how DSB affect the separation of two loci, we computed the MFDT for untreated and Zeocin treated cells (Fig. 3C) that shows an increase pattern with *∊*. The MFDT for the untreated case increased from 0.2 to 0.9 s, whereas the MFDT for Zeocin treated increased from 0.2 to 0.5 s as *∊* increased from 0.15 to 0.5 *μ*m. We conclude that it takes less time for the two loci to dissociate following DSB, probably due to the chromatin decompaction, enhanced mobility and the activity of repair proteins. This result suggests that repair molecules do not impair the local chromatin motion.

We conclude that uniformly distributed DSBs impair the MFET only at a scale below 0.25 *μ*m, suggesting that this scale characterizes the local chromatin organization in which undamaged loci can freely move, but become restricted above it. These finding confirms the confinement found in [1] (Supplementary Fig.5b), which reported that a radius of confinement of 0.23 *μ*m for Zeocin 500 *μg/ml* treatment. However, we show here that following DSBs, the local exploration of the chromatin remain characterized by recurrent motion and if repair molecules do affect the encounter time at a spatial scale below 0.25 *µ*m, they do not prevent the dissociation time of the two loci.

### 3.3 Stochastic simulations of a DSB in randomly cross-linked (RCL) polymer

To further investigate the difference between chromatin reorganization before and after DSBs, reported above for the two loci dynamics, we now use a Randomly Cross-Linked (RCL) polymer, generalizing the Rouse polymer model (Methods), to evaluate the changes in the constraints of SPTs statistics following DSBs. The parameters of the RCL model are calibrated from experimental data [1], in which two tagged monomers are tracked before and after DSB induction between the tagged loci. The tagged loci were tracked over 60 s at a time interval of 300 ms. The confinement length [25] is defined as

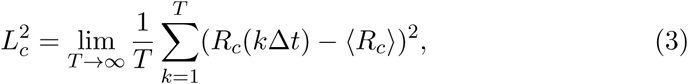

and we computed it before and after DSB induction, where *R*_*c*_(*t*) is the vector position between the two tagged loci at time *t*. As reported in [2], for an unbroken locus *L*_*c*_ = 0.13 *μ*m, and *L*_*c*_ = 0.23 *μ*m after DSB induction.

To identify the possible local chromatin reorganization underlying this difference in *L*_*c*_, we simulate a RCL polymer (Methods) with *N* = 100 monomers, containing an additional *N*_*c*_ connectors between randomly chosen non-nearest neighboring monomers [13, 18, 19] (Fig. 4A). The added connectors reflects the compaction in the coarse-grained representation of the chromatin by molecules such as cohesin CTCF and condensin [26]. Randomly positioning connectors reflects the heterogeneity in chromatin architecture in a population of cells.

**Figure 4:**
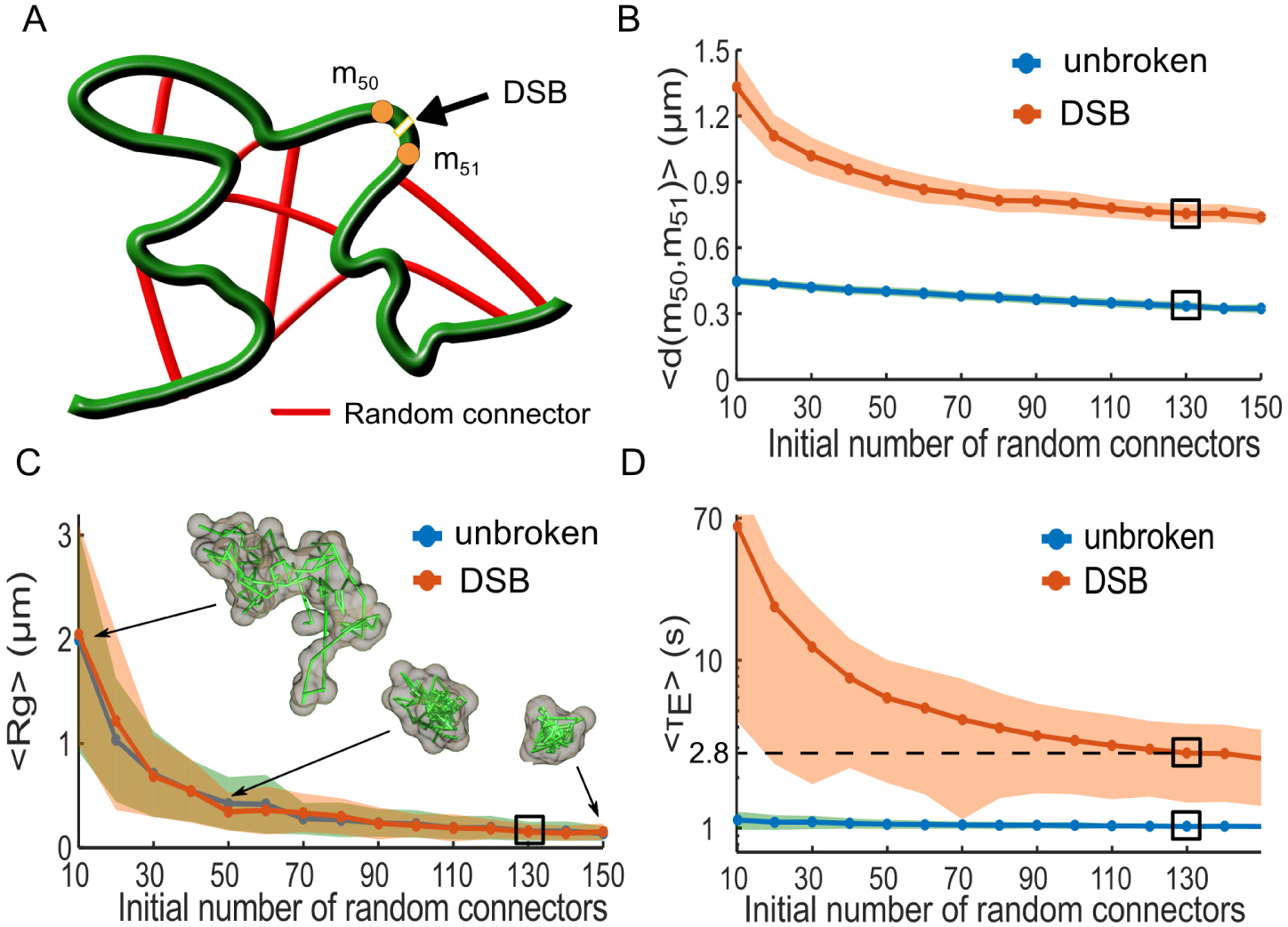
Local force destabilization following a double-strand DNA break (DSB). **A.** Schematic representation of a randomly cross-linked (RCL) polymer, where *N*_*c*_ random connectors (red) are initially added to the linear backbone (green) of a Rouse chain. A DSB is induced between monomers *m*_50_ and *m*_51_, modeled by removing the spring connectors between them and all random connectors to these monomers. **B.** Mean maximal distance 〈*Max*(*d*(*m*_50_,*m*_51_))〉 for both the unbroken loci (blue) and DSB (orange) simulations, where the shaded are the STD. The black rectangle indicates the value obtained for *N*_*c*_ = 130 matching *L*_*c*_ (eq. 3) measurements reported in [2], where we obtain 0.37 *μ*m for the unbroken and 0.86*μ*m for DSB simulation **C.** The mean radius of gyration (MRG), 〈*R*_*g*_〉, obtained from simulations of 100 monomer RCL polymer (blue) and after DSBbetween monomers 50 and 51 (orange). Three sample polymer realizations are shown for *N*_*c*_ = 10, 50, and 150. For *N*_*c*_ = 130 we obtain 〈*R*_*g*_〉 = 0.15*μ*m for both cases **D.** The mean first encounter time (MFET) 〈*τ*_*E*_〉 for *m*_50_ and *m*_51_ plotted with respected to *N*_*c*_ for both the unbroken (blue) and DSB(orange) simulations. The MFET is displayed on a semi-log axes, where before DSB we obtained 〈*τ*_*E*_〉 and 2.8 s following DSB and the removal of 5 random connectors on average.

We first find the minimal number of random added connectors *N*_*c*_ by varying *N*_*c*_ in the range 10-150 for both unbroken loci and after DSB induction. We then computed *L*_*c*_ from simulations (1000 realizations for each *N*_*c*_) and adjusted *N*_*c*_ to match the measured one. For each realization, we randomized the choice of monomer pairs to connect. We simulated each realization until the relaxation time *τ*_*R*_ = *b*^2^*D/*(3*λ*_1_), where *b* is the standard-deviation of the vector between adjacent monomers, and *λ*_1_ is the smallest non-vanishing eigenvalue of the polymer’s connectivity matrix [27], which we calculated numerically. We empirically found *λ*_1_ to vary between 0.15 when *N*_*c*_ = 10 to 0.8 for *N*_*c*_ = 150, resulting in *τ*_*R*_ in the range of 20 minutes to 23 s until polymer relaxation. We then continued the simulations for an additional 200 steps at Δ*t* = 300 ms for a total of 60 s, to match the experimental recorded time [1]. For DSB simulations, we induced a DSB between monomer 50 and 51 after the relaxation time *τ*_*R*_ and we then removed the spring connector between them. To account for the local chromatin decompaction, we further removed all random connectors to monomers 50 and 51. We discarded polymer configurations where the polymer chain was divided into two separated chains after the induction of DSB and removal random connector. Simulation parameters are summarized in Table 1, where *N*_*c*_ remains a free parameter.

We computed the average values of *L*_*c*_ for monomers 50 and 51 over each realization and found a good agreement between simulations and experimental data when *N*_*c*_ = 130 for both the unbroken (*L*_*c*_ = 0.13 *μ*m) and broken loci (*L*_*c*_ = 0.23 *μ*m). After DSB induction, we recover the value *L*_*c*_ = 0.23 *μ*m for *N*_*c*_ = 125, where 5 connectors, on average are removed between. Furthermore, the mean maximal distance between monomers 50 and 51 (Fig. 4B) decayed from 0.4 to 0.3 *μ*m when *N*_*c*_ varied between 10 and 150 for the unbroken loci simulation, while it changes from 1.3 *μ*m for *N*_*c*_ = 10 to 0.75 *μ*m for *N*_*c*_ = 150 in the DSB simulations. When *N*_*c*_ = 130 the mean maximum monomer distance was 0.33 *μ*m and increased to 0.75 *μ*m after DSB induction.

Using the mean radius of gyration 〈*R*_*g*_〉 (Fig. 4C) computed for RCL polymer configuration, we show that compaction increases with *N*_*c*_. Thus〈*R*_*g*_〉 decreased from 2 *μ*m for *N*_*c*_ = 10 to 0.15 *μ*m at *N*_*c*_ = 150. Note that DSBs do not affect the radius of gyration 〈*R*_*g*_〉 for all *N*_*c*_ ∈ [10, 150] (Fig. 4C). In that figure, we further represented three polymer realizations for *N*_*c*_ = 10, 50, 150. For *N*_*c*_ = 130, the value of the gyration radius is 〈*R*_*g*_〉 = 150 nm for both the unbroken and DSB (numerical simulations).

To further examine the relationship between the chromatin local architecture and transient properties of the chromatin, we computed the first passage time (FET) between monomer *m*_50_ and *m*_51_, before and after DSB induction (Fig. 4A). Each simulation realization was terminated when *m*_50_ and *m*_51_ enter for the first time within a distance less than ∊, where we recorded the first encounter time *τ*_*E*_. Terminating each simulation after the first encounter allowed us to randomize the position of connectors for any other simulation and thus better account for chromatin structural heterogeneity. The MFET for the unbroken loci simulations decreases from 1.1 s to 1 s when *N*_*c*_ increases from 10 to 150, while it decreases from 62 s for *N*_*c*_ = 10 to 2.6 s for *N*_*c*_ = 150 (Fig. 4C) following DSB. Interestingly, when *N*_*c*_ = 130, only 5 connectors were removed on average to account for a DSB, but before induction 〈*τ*_*E*_〉 between *m*_50_ and *m*_51_ was 1 s and increased to 2.8 s after DSB induction. These time scales are consistent with data used in figures 1 and 2.

We conclude that the empirical confinement length can be accounted for in RCL polymer using a *N*_*c*_ = 130 connectors. Following a DSB the average number of removed connectors was 5, which represents 4% of the total number of connectors *N*_*c*_. Interestingly, the mean radius of gyration 〈*R*_*g*_〉 ≈ 150 nm is mostly unchanged between the unbroken and DSB. However, the encounter time *τ*_*E*_ (Fig. 4D changed from 1 to 2.8 s, showing that removing the key connector could affect the local encounter time. The drastic effect of changing the number of connectors appears in the mean maximal distance between the two monomers, increasing from 0.33 *μ*m in the unbroken case to 0.75 *μ*m after DSB, leading to high increase in the local search time. This search in local restricted environment could be at the basis of the Non-Homologous end joining (see Discussion).

## 4 Discussion

We introduced here a transient analysis of loci trajectories based on computing the first encounter times between two simultaneously tagged chromatin loci to a small distance. Because the positions of the loci fluctuate in time but return recurrently into close proximity, this dynamics generates enough statistical events. We showed here that this statistics revealed a characteristics length around 200 nm, where the chromatin constrains the two loci dynamics. This analysis cannot be obtained from the traditional parameters, extracted from SPTs such as the mean square displacement (MSD) or the anomalous exponent [20], which characterize the dynamics of individual locus separately. Such parameters were used in the past to study the deviation from Brownian motion [21]. Further information about chromatin organization is obtained from individual single loci trajectories [22], such as the length of constraint characterizing confinement or the tethering force to account for the first statistical moment and the mean force responsible for confinement [6, 9, 10, 2, 1, 28].

The statistics of the FET and FDT account for the correlated properties of two loci and are directly related to the transient properties of the chromatin: the FET reveals that the recurrent visit time between the loci, depending on the genomic distance (Fig. 1), varies from 1.5 to 2.5 s. We further confirm that altering the chromatin integrity by generating DNA damages using the Zeocin drug [1] affects the MFET at a distance lower than 200 nm (Fig. 2B), showing that this scale is certainly critical in chromatin remodeling [1]. Above this distance, the MFET was constant and we interpret this result as a consequence of the local crowding effect. These results further show that the recurrent visits between two loci can be modulated by chromatin remodeling. The confinement length of few hundreds of nanometers estimated here is compatible with the one extracted from Hi-C data of the order of 220 nm, using polymer looping in confined microdomains [15]

To further explore how chromatin re-organization affects recurrent loci encounters, we use the RCL polymer model [19], which is a Rouse polymer with added random connectors. This approach consists in adding random connectors is more accurate in representing tethering force than the average computation that we introduced in [10]. Random cross-linking in the RCL model serves to simulate the confined environment [29] and the heterogeneity in chromatin architecture in cell population. We used the changes in the length of constraint reported in [1] to calibrate the number of added random connectors and simulated trajectories of the RCL before and after the induction of DSB. Interestingly, the consequence of DSB damages on chromatin reorganization is equivalent of removing 4% of the connectors in the vicinity of the DSB, leading to an increase of distance between the two broken part from 0.4 to 0.9 *μ*m, while the mean radius of gyration 〈*R*_*g*_〉 was almost unchanged at 0.15 *μ*m. However, the MFET increased from 1 to 2.8 s. The random cross-links in the RCL model thus play the role of the confining environment, which prevents the two ends from drifting apart (Fig 4B and C), similarly to the crowding effect seen in [29] for self-avoiding polymers. Bending elasticity and self avoidance could be accounted for by altering the number of random connectors.

The present model reveals that the local confined decompaction following DSB prevents the two ends to drift apart, which could have drastic consequences in dsDNA break repair processes, such as during non-homologous-end-joining (NHEJ), where the two ends should be re-ligated together. The possible role of stabilizing the broken ends by maintaining a large number of connectors is probably to avoid inappropriate NHEJ religations that can lead to translocations or telomere fusion. We remark that the MFET that we computed here cannot be used to study the other repair process called homologous recombination, which is based on a long-range spatio-temporal search for a homologous template [30, 6]. We conclude that the present first passage time statistics derived from polymer simulations can be used to analyze any temporal correlation between loci pairs. It would certainly be interesting to record three loci simultaneously at different distances and apply our method to it to obtain refine properties of chromatin reorganization.

## Funding

This work was supported by the Marie-Curie award and Labex MemoLife.

## Acknowledgments

We thank Tom Owen-Hughes lab for sending us their SPTs data. Data sets were previously published in [1, 22]. DH thanks the Simons foundation for support. This research was supported by a Marie Curie Award to DH.

## 5 Theory and Methods

### 5.1 Looping times in chromatin polymer models

To analyze the statistics of two loci located on the same chromatin arm, we use the classical Rouse polymer model that describes a collection of beads *R*_*n*_(*n* = 1*…N*) connected by harmonic springs and driven by Brownian motion [13]. The energy of the polymer is given by

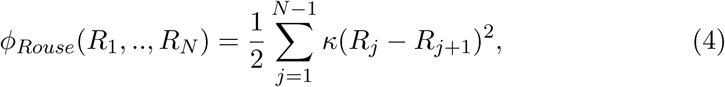

where 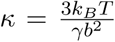 is the spring constant, *b* is the standard deviation of the connector between adjacent monomers, *γ* is the friction coefficient, *k*_*B*_ the Boltzmann coefficient, and *T* the temperature.

The first encounter time (FET) between two loci is defined as the first time the two loci are positioned within a ball of radius *∊*. The distribution of FET between the two ends of a polymer chain is well approximated by a Poisson process in free and confined domains [15, 23]. In both cases, the distribution of the decay rate constant *λ*_*E*_ is the reciprocal of the mean first encounter time (MFET) 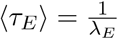 and the probability density function is

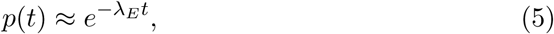

In a confined domain, the expression for the MFET is

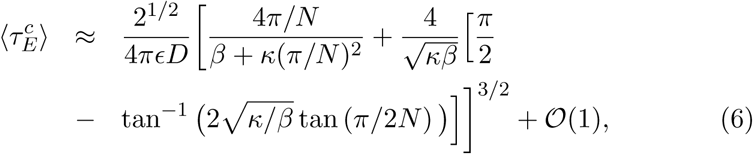

where

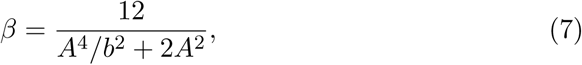

and *A* is the radius of a sphere confining the polymer [15].

### 5.2 Dissociation times in a parabolic potential

To characterize the dissociation time of two loci, we adopt the Kramer’s escape over a potential barrier [31]. The potential can be due to the average forces between local monomers. We model it as an effective parabolic well truncated at a hight *H*. In the deep circular well approximation of size *a*_0_ [31], the escape time for a process 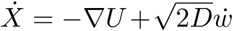 is (in two dimensions)

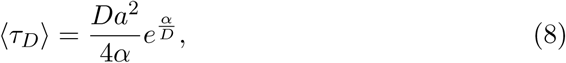

where 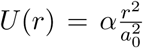 and the energy is *E* = *U* (*a*) = *α* and *U* (0) = 0. The distribution of escape time is Poissonian with rate 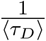.

For the effective problem of unlooping to a certain distance, we consider that this problem is equivalent to the escape of a particle from a well with diffusion coefficient *N D*, where *N* is the number of monomers. In the present case, *N* is proportional to Δ and we have used the empirical formula:

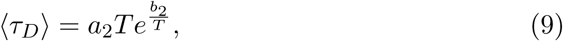

where *a*_2_ and *b*_2_ are two constants.

### 5.3 Construction of the randomly cross-linked (RCL) polymer model

The Rouse polymer [13] describes chromatin below a scale of few Mbp [32, 16]. Starting from a Rouse model [13], the RCL is constructed by adding sparse connected pairs (Fig. 4A red), chosen with a uniform probability such that the potential for the polymer is the sum of the *ϕ*_*Rouse*_ plus the potential

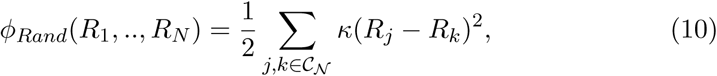

where C_N_ is an ensemble of indices from 1 to *N*. The chromatin is modeled as a polymer chain with a uniform variance *b*^2^ between adjacent monomers.

The total energy of a polymer containing random connectors is the sum of two energies 4, and 10

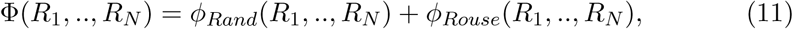

and the stochastic equation of motion for *n* = 1, .., *N* is

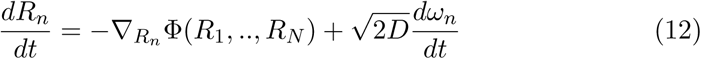

where 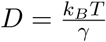is the diffusion constant, *γ* is the friction coefficient, and *ω*_*n*_ are independent 3-dimensional Gaussian noise with mean 0 and standard deviation 1. We use this construction to estimate the minimal number of connectors before and after a dsDNA-breaks.

Simulations of the RCL polymer were performed using codes written in Julia v0.5.1 [33]. Codes are available on the Bionewmetric website http://bionewmetrics.org/. We summarize in Table 1 the values of parameters used in simulations

